# Type 2 diabetes increases susceptibility to invasive *Salmonella* Typhimurium despite butyrate supplementation

**DOI:** 10.64898/2026.08.14.744872

**Authors:** Cecilia G. Sierra-Bakhshi, Laura A. Farr, Michael E. Smith, Torren A. Kalaskey, Katelyn G. Perkins, Maria G. Winter, Saroj Sigdel, Sebastian E. Winter, Lydia M. Bogomolnaya

**Affiliations:** Department of Biomedical Sciences, Joan C. Edwards School of Medicine, Marshall University, Huntington, WV; Department of Internal Medicine: Division of Infectious Diseases, UC Davis, Davis, CA; Department of Pathology, Joan C. Edwards School of Medicine, Marshall University, Huntington, WV; Department of Medical Microbiology and Immunology, UC Davis, Davis, CA; Center for Immunology and Infectious Diseases, UC Davis, Davis, CA

## Abstract

Non-typhoidal *Salmonella* is a major cause of bacterial foodborne illness leading to acute gastroenteritis. In individuals with type 2 diabetes (T2D), *Salmonella* infection is more likely to cause life-threatening extraintestinal infections. The mechanism underlying this susceptibility remains unclear. In this study, 8-week-old TALLYHO mice were fed either a chow or high-fat diet (HFD, 45% fat) for 8 weeks to induce the T2D. As expected, HFD-fed mice gained more weight and developed diabetic-range blood glucose levels by 16 weeks of age. Next, mice from each diet group were orally infected with a fully virulent bioluminescent *Salmonella* Typhimurium to monitor infection spread by *in-vivo* imaging. Although both groups developed clinical signs of salmonellosis, *Salmonella* spread was accelerated and followed an unusual pattern in T2D mice compared with healthy animals. Additionally, hyperglycemia increased gut-derived lipopolysaccharide leakage into the bloodstream. Based on the link between T2D and altered levels of butyrate-producing bacteria in the gut, we analyzed the intestinal short-chain fatty acid (SCFA) profiles in the TALLYHO mice. As expected, intestinal SCFA concentrations, including butyrate, were lower in HFD mice than in chow-fed animals. Given butyrate’s role in gut health and its ability to downregulate *Salmonella* invasion genes, mice received oral butyrate supplementation. We found that butyrate supplementation reduced the extraintestinal spread of *Salmonella* in normoglycemic chow-fed animals. Unexpectedly, although butyrate improved intestinal health in hyperglycemic mice, it failed to decrease *Salmonella* spread in diabetic animals. Taken together, these findings provide novel insights into the pathogenesis of enteric salmonellosis in the context of T2D.

## Introduction

Type 2 diabetes mellitus (T2D) is a metabolic condition primarily defined by the development of insulin resistance and decreased pancreatic β-cell function that consequently results in chronic hyperglycemia. According to the report from the Global Burden of Diseases, Injuries, and Risk Factors Study (GBD), 529 million people were living with diabetes in 2021, and 96% of those individuals have type 2 diabetes (1). Research estimates predict that by 2030, diabetes will become the seventh leading cause of death with 3% of all deaths globally due to the disease (2). Furthermore, by 2050, more than 1.31 billion people are projected to have diabetes (1). Obesity along with genetics, lifestyle, and diet are recognized as the primary factors in the development of type 2 diabetes, and the combined adverse health effects of both conditions are known as diabesity (3). The GBD study showed that 2.1 billion adults were overweight or obese in 2021 and forecast that over half of the global adult population will be impacted by these conditions by 2050 (4). The increase in obesity rate and in T2D incidence are fueled by calory-rich high fat “westernized” diet and sedentary lifestyle typical of modern society (5, 6). Consumption of a high-fat diet in combination with lack of exercise results in a state of energy imbalance leading to hyperglycemia requiring higher circulating levels of insulin.

People with diabetes are more vulnerable to infections, and these infections are often more severe compared to the general population (7). For instance, in individuals with T2D, *Salmonella* infections are three times more likely to occur (8). Additionally, the risk of extraintestinal *Salmonella* infections also increases in diabetic patients, leading to higher mortality rates (9, 10). Moreover, T2D-associated salmonellosis often lacks the initial gastrointestinal symptoms and leads to life-threatening conditions including but not limited to suppurative thyroiditis, soft tissue infections, osteomyelitis, pyomyositis, myonecrosis and pericarditis (11–22). Although the increased susceptibility of T2D patients to *Salmonella* was originally linked to reduced acidity in the stomach and impaired intestinal motility (23), hyperglycemia has also been reported to cause damage to the intestinal epithelial layer (24–26). Despite the relevance of this phenomenon to human health, the underlying mechanism in the extraintestinal spread of *Salmonella* in the context of diabetes is poorly understood.

A key contributing factor to the limited research on this topic is the lack of an appropriate mouse model of human type 2 diabetes, since the traditional models used to research diabetes do not accurately reflect the natural process of development of this chronic disease. For instance, *ob/ob* and *db/db* mice are commonly used in T2D research; however, these animals carry autosomal mutations in the leptin gene (*ob/ob*) or its receptor (*db/db*) leading to disruption in leptin signaling (27). As a result, these mice develop hyperphagia, reduced energy expenditure, obesity and insulin resistance at an early age (28). Because human T2D generally lacks severe juvenile-onset obesity and insulin resistance, concerns of translational relevance of findings derived from monogenic T2D models were raised (29). Another common practice is to utilize nutritional models of T2D. One such model employs C57Bl/6 mice fed a high fat diet (40-60%) for 6 months, leading to obesity and diabetes (30). Although C57BL/6J mice develop moderate hyperglycemia, a shortcoming of this model is the length of time required to induce hyperglycemia development, which can be up to six months of consuming a high fat diet (HFD) (31). The polygenic rodent models of T2D, including TALLYHO/Jng mouse strain, overcome many of these limitations and provide an opportunity to study diabetic phenotypes that are more consistent with human profiles (28). TALLYHO (TH) mice have normal leptin and leptin receptor genes and, therefore, develop less severe metabolic syndrome compared to more popular diabetic *db*/*db* mice with monogenic obesity (32). TH mice display normal levels of glucose in the blood at weaning, but the levels spontaneously rise with age in males. Male TH mice develop persistent hyperglycemia, insulin resistance, hyperinsulinemia, and hyperlipidemia around 14-16 weeks of age (33). This phenotype is not 100 percent penetrant (34) but it can be promoted by diet with high fat content (35).

Type 2 diabetes is a progressive metabolic disease that has a negative impact on multiple organ systems including changes in the intestines and in the composition and distribution of gut microbiota. Several metagenomic studies highlighted the reduction of butyrate-producing bacteria in stool samples of diabetic patients compared to healthy individuals (36–40). Butyrate, a short chain fatty acid exclusively produced in the lower intestinal tract through the fermentation of dietary fiber, has been found to have many beneficial properties. Butyrate is not only a major energy source for colonic epithelial cells, but it has also been found to reduce mucosal inflammation, oxidative status, and reinforce the intestinal epithelial barrier (41–43). Butyrate has been shown to be able to reduce body weight, modulate and lower blood glucose levels including HbA1C measurements, insulin resistance, fat accumulation, dyslipidemia, and diabetic inflammation (44–46). Besides direct benefits to the host, butyrate is also known to have antimicrobial properties against several bacteria species including *Acinetobacter baumanii*, *Staphylococcus pseudintermedius*, *Clostridium perfringens* and *Salmonella* Typhimurium (47–49). However, the ability of butyrate to limit *Salmonella* colonization in the context of diabetes was not previously addressed.

Here, we utilize TALLYHO/Jng mice to study *Salmonella* colonization in the diabetic host and show that chronic hyperglycemia accelerates extraintestinal spread of the pathogen. We also show that blood lipopolysaccharide (LPS) levels are elevated in diabetic animals indicating increased gut permeability. Furthermore, the intestinal butyrate levels are reduced in hyperglycemic animals. Finally, we show that butyrate supplementation significantly reduces extraintestinal spread of *Salmonella* in normoglycemic animals. However, while butyrate supplementation improves intestinal health in diabetic mice, it fails to limit extraintestinal spread of *Salmonella* in hyperglycemic animals.

## Results

### TALLYHO mice develop persistent hyperglycemia by 16 weeks of age

To promote diabetes development, we divided 8-week-old male TH mice into two groups. The average body weight of mice at this age was 30.67±2.76 grams. One group was maintained on a chow diet throughout the study, while the other group was switched to the 45% high fat diet (HFD). We monitored weight gain weekly until animals reached 16 weeks of age (Figure 1A). Chow-fed mice continued to grow until 14 weeks of age. At this point chow-fed animals gained 4.45±1.83 grams. Mice in HFD group gained weight significantly faster compared to the control group. By 14 weeks of age the average weight gain for this group was 7.86±4.12 grams.

**Figure 1.**
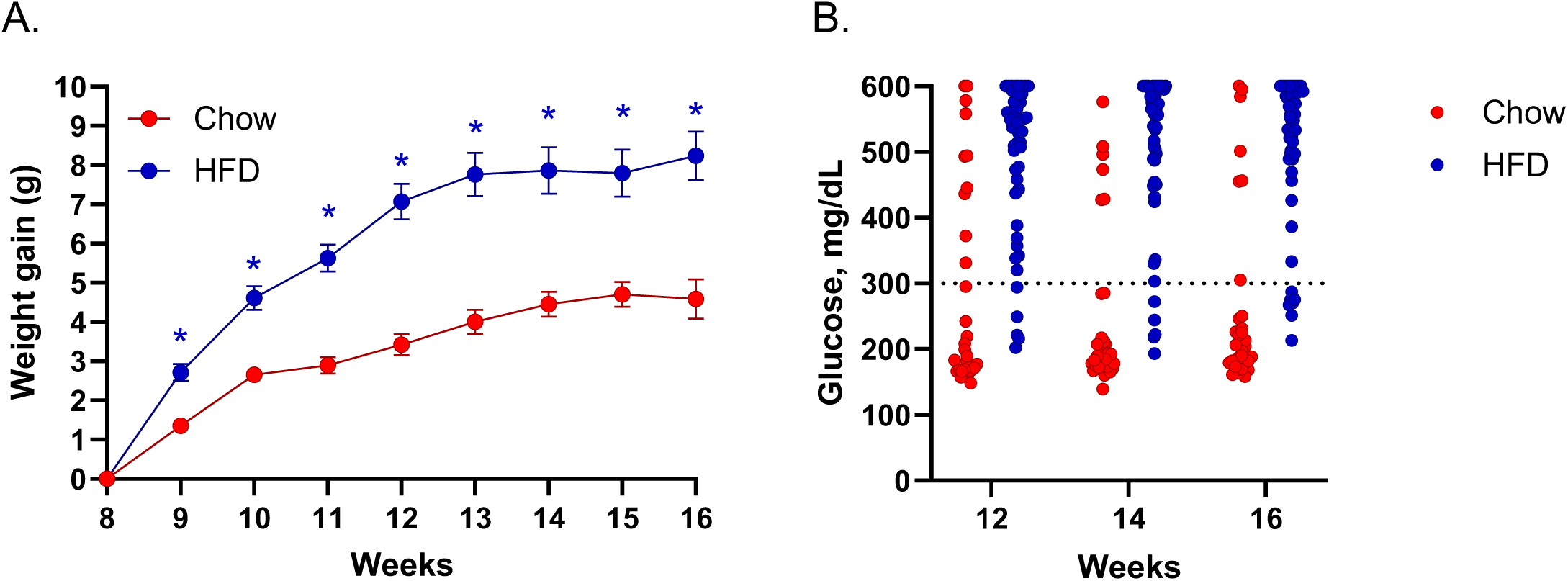
High fat diet promotes weight gain and hyperglycemia in male TALLYHO mice. **A.** Weight gain was followed weekly. **B.** Non-fasting blood glucose was measured using OneTouch Ultra 2 glucometer. Mice were considered hyperglycemic when the average glucose concentration was above 300 mg/dL. Asterisk indicates significance in multiple unpaired t-test with Holm-Šìdák method, *p*<0.05

Alongside weight, we also monitored non-fasting blood glucose level in animals on both diets. Most chow-fed mice stayed normoglycemic, though approximately 18% of animals became hyperglycemic (Figure 1B). In contrast, 86% of animals from HFD group developed persistent hyperglycemia.

In chow-fed mice chronic hyperglycemia resulted in significant enlargement of spleen, liver, and kidney; however, in HFD-fed animals, chronic hyperglycemia led to significant enlargement of liver and kidney, but not the spleen (Figure S1).

Mice with and without hyperglycemia (>300 mg/dL non-fasting blood glucose) in HFD group and animals with a normal level of blood glucose in the chow group were selected for the infection study. Overall, these findings suggest that TH mice fed a high-fat diet may be suitable model to assess the impact of hyperglycemia and obesity on infections with enteric pathogens.

### Type 2 diabetes promotes systemic spread of *Salmonella* in TALLYHO mice

Commonly used strains of inbred mice respond to *Salmonella* exposure differently: BALB/c, C57BL/6 are highly susceptible to bacterial infection due to mutated *Slc11a1* allele, while A/J, CBA/J, and 129 sv mice can effectively limit colonization of organs outside of the intestines (50–52). Susceptibility to infection in TH mice was not previously investigated. To establish the infection outcome in these animals, 16-week-old chow-fed normoglycemic male TH mice were inoculated intragastrically with 1x10^6^ CFU of fully virulent bioluminescent *Salmonella* Typhimurium (Figure 2A). Progression of infection was monitored by tracking the bioluminescence spread *in vivo*. The detectable signal appeared on day 4 post infection (Figure 2B), and by day 5 the bacteria were observed throughout the body (Figure S3). Interestingly, infection of HFD-fed normoglycemic TH mice with *Salmonella* resulted in a similar colonization dynamic to the chow-fed animals. In contrast, in hyperglycemic HFD TH mice, the bioluminescent signal was detected as early as day 2 post infection with atypical extraintestinal localization (Figure 2B). Quantification of luminescence confirmed that the signal intensity was similar between chow and HFD mice with normal blood glucose levels. However, the intensity of luminescence was significantly higher in HFD mice with hyperglycemia starting from day 2 and continuing through day 4 post infection compared to chow-fed normoglycemic animals (Figure 2C). Furthermore, HFD animals with hyperglycemia developed signs of infection significantly sooner compared to the control chow mice and had to be euthanized on day 4 post infection (Figure 2D).

**Figure 2.**
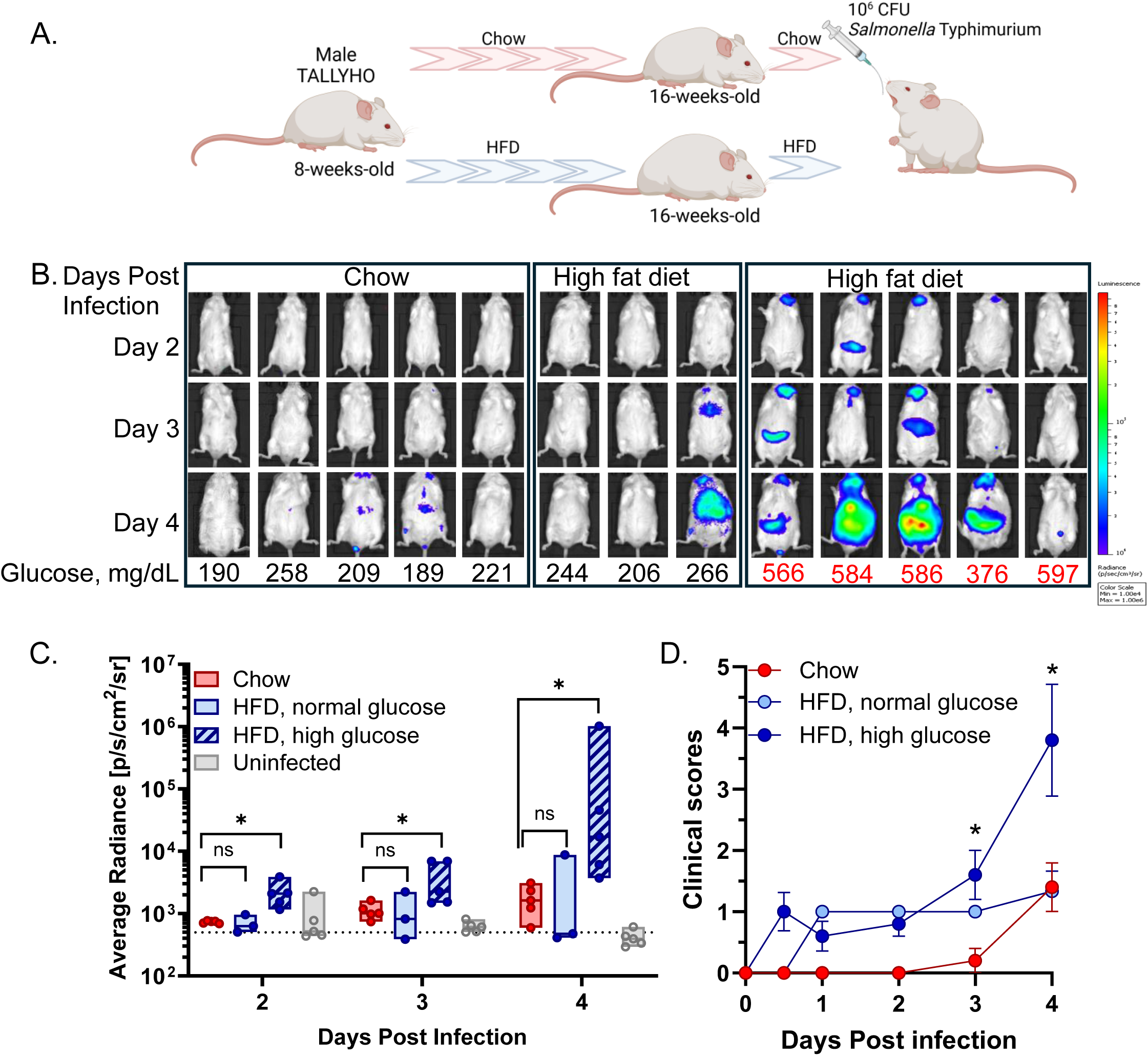
Hyperglycemia promotes susceptibility to *Salmonella* infection in male TALLYHO mice. **A.** Experimental design of pre-infection TH mice treatments. Created with BioRender.com. **B.** 16-weeks-old male TH mice were orally infected with 1x10^6^ CFU of *Salmonella* strain expressing chromosomally encoded *hisG::rpsM-luxCDABE* and imaged using IVIS Lumina XRMS III system. **C.** Bioluminescence quantification was done using Living Image software. Asterisk indicates significance in multiple Asterisk indicates significance in multiple Mann-Whitney U test with Holm-Šìdák method, *p*<0.05. **D.** Clinical scores post infection. Asterisk indicates significance in two-way ANOVA with Šìdák’s multiple comparison’s test.

### Hyperglycemia impairs gut barrier function in TALLYHO mice

We hypothesized that the increase in extraintestinal *Salmonella* spread in hyperglycemic HFD TH mice (Figure 2B) was at least in part due to intestinal barrier disruption in the diabetic animals. The increase in lipopolysaccharide (LPS) translocation from the intestine to the bloodstream was previously detected in T2D patients (53). Additionally, a high fat diet further promotes impairment of the gut barrier function through disruption of the tight junction proteins in the intestines (54).

To test if TALLYHO mice recapitulate this aspect of type 2 diabetes previously noticed in the monogenic rodent models of disease (26), we measured LPS concentration in the serum of TH mice with and without hyperglycemia from chow and HFD groups (Figure 3A). We found that the circulating LPS levels were indeed significantly higher in hyperglycemic obese HFD mice compared not only to the control chow-fed animals but also to the normoglycemic HFD-fed mice.

**Figure 3.**
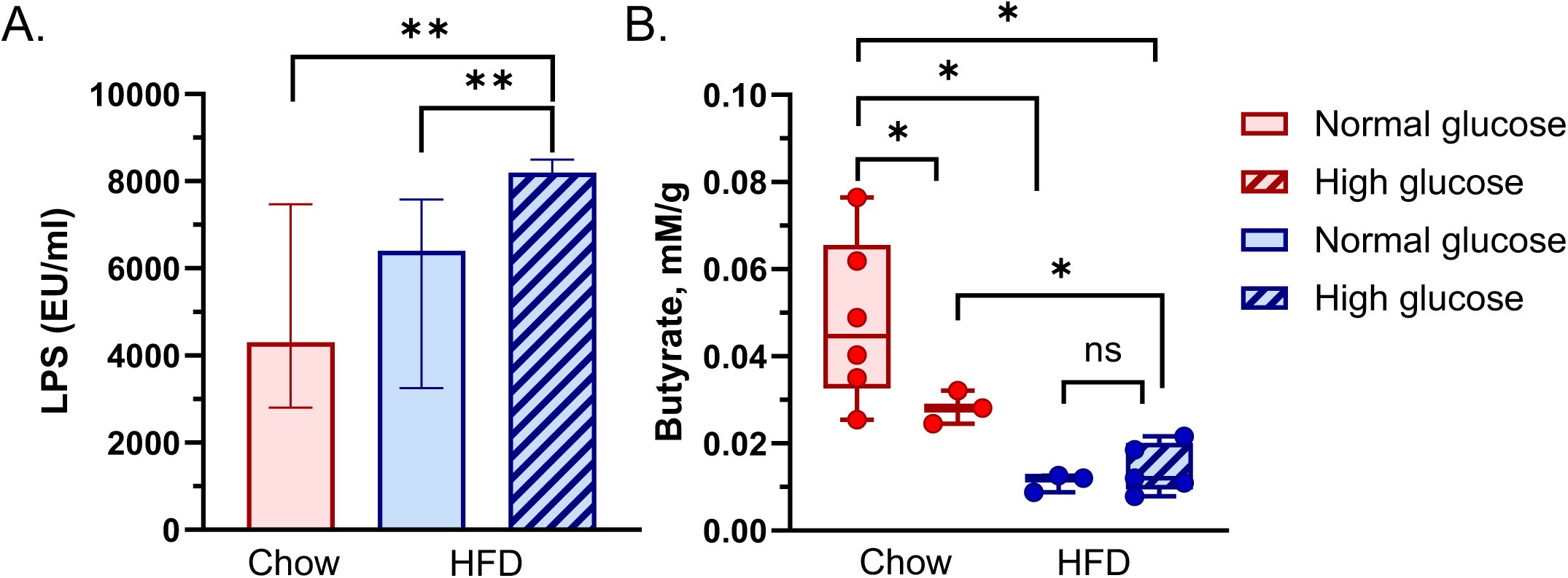
Hyperglycemia promotes increased intestinal permeability and leads to reduction in cecal butyrate in male TALLYHO mice. **A.** Serum LPS levels were measured using PyroGene recombinant factor C endpoint fluorescent assay (Lonza) and shown as median with interquartile range. Asterisk indicates significance in multiple Mann-Whitney U test with Holm-Šìdák method, *p*<0.05. **B.** Cecal butyrate concentration was determined by GC-MS using multiple reaction monitoring mode. Asterisk indicates significance in multiple unpaired t test with Welch correction test using Holm-Šìdák method, *p*<0.05

### Intestinal butyrate is reduced in diabetic TALLYHO mice

Recent metagenomics studies have shown that type 2 diabetes patients have altered gut microbiota with a decreased population of butyrate-producing bacteria (36–40). To test whether butyrate concentration is altered in TH mice with diabetes, we collected cecal content from TH mice with and without hyperglycemia from chow-fed and HFD groups and measured short chain fatty acid (SCFA) concentrations in the samples using gas chromatography-mass spectrometry (GC-MS) (Figure S2). We found that the total SCFA concentrations, including butyrate, were lower in chow-fed hyperglycemic mice than in normoglycemic animals on the same diet (Figures S2 and 3B) indicating that type 2 diabetes development in TH mice accurately reproduces this aspect of human disease. Butyrate concentration was further reduced in the cecal content of animals from HFD group regardless of glucose levels in their blood (Figure 3B).

Butyrate is a major molecule used as an energy source by colonocytes, but it has also been suggested to reduce inflammatory responses in the colon and to help maintain the intestinal barrier (42, 55–57). In order to test whether butyrate supplementation can aid in restoration of gut integrity, we treated normoglycemic chow-fed mice and hyperglycemic HFD animals with a daily oral dose of tributyrin (butyrate prodrug) for five consecutive days (Figure 4A). As expected, butyrate supplementation resulted in a significant reduction of blood LPS in chow-fed mice (Figure 4B). More importantly, butyrate treatment also reduced the amount of LPS in circulation in hyperglycemic HFD animals to the levels comparable to the one present in the chow-fed normoglycemic mice (Figure 4B).

**Figure 4.**
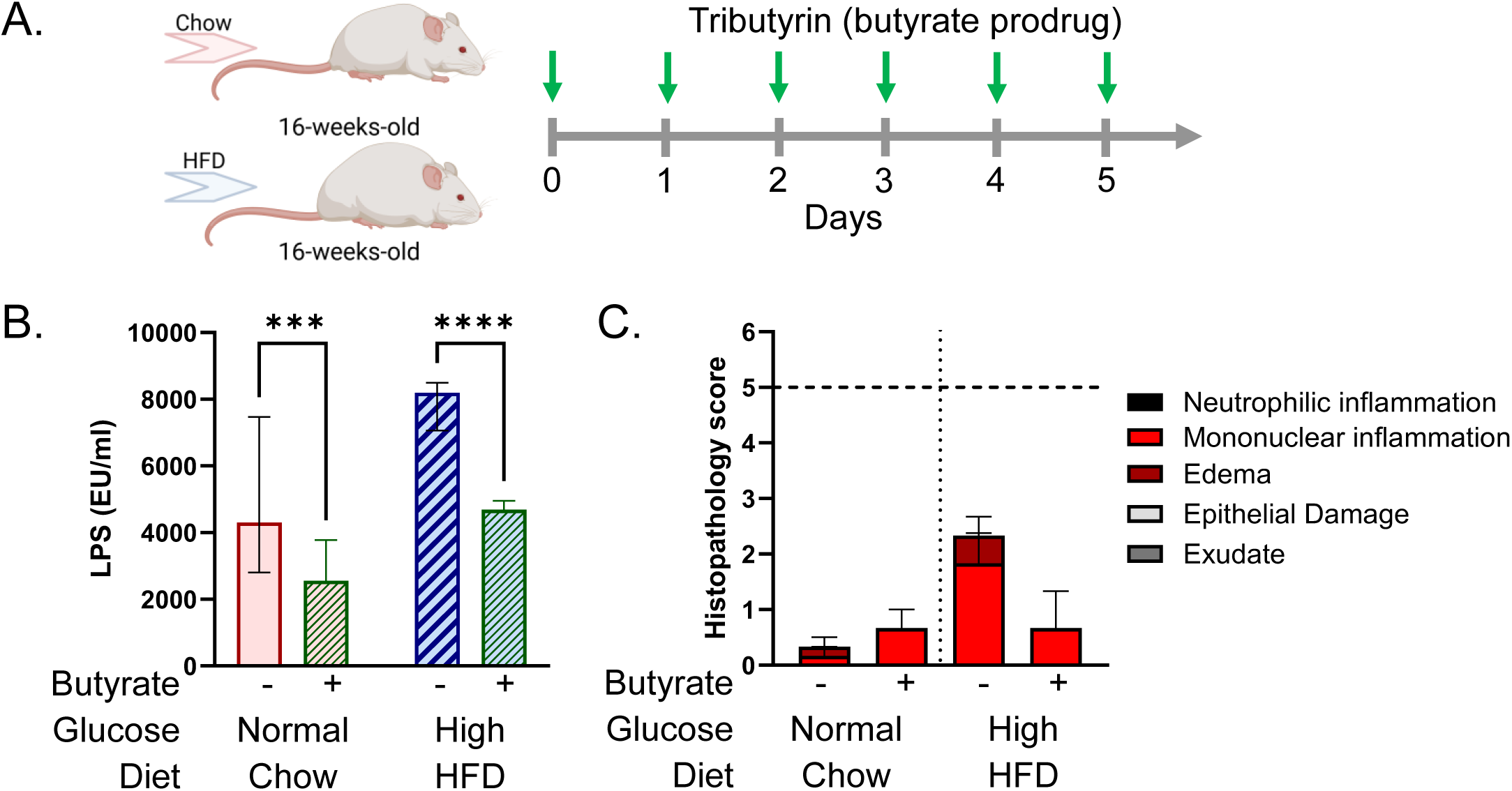
Butyrate improves intestinal health in diabetic mice. **A.** Schematics of tributyrin treatment. Created with BioRender.com. **B.** Butyrate supplementation reduces LPS leakage from the intestines. Serum LPS levels were measured using PyroGene recombinant factor C endpoint fluorescent assay (Lonza) and shown as median with interquartile range. Asterisk indicates significance in multiple unpaired t test with Welch correction test using Holm-Šìdák method, *p*<0.05. **C**. Sections of colon were collected from 16-week-old TH mice, fixed in formalin, paraffin embedded, cut, and stained with hematoxylin and eosin. Stained sections were scored for the signs of inflammation. A combined score of <5 corresponds to normal or mild inflammation; a combined score of >5 indicates moderate to severe inflammation.

Consumption of a diet rich in fat leads to low-grade intestinal inflammation (58). Accordingly, we noticed a slight increase in histopathology score in the colon of HFD TH mice compared to the chow-fed animals (Figure 4C). In agreement with previous observations (59), butyrate treatment reduced inflammation in the HFD animals (Figure 4C).

Because butyrate is also known to limit the luminal expansion and to downregulate epithelial invasion of *Salmonella* (60–62), we hypothesized that oral supplementation with tributyrin might limit the extraintestinal spread of the pathogen in TH mice.

### Butyrate limits extraintestinal spread of *Salmonella* in healthy but not in diabetic TALLYHO mice

To test if butyrate supplementation can prevent the severity of *Salmonella* infection, we began our study by orally infecting normoglycemic chow-fed mice with 1x10^6^ CFU of bioluminescent *S*. Typhimurium (STM). These animals were immediately treated with a daily oral dose of tributyrin for five consecutive days. Butyrate supplementation protected mice from severe weight loss following STM infection (Figure 5A) and reduced the overall clinical scores (Figure 5B) compared to the untreated infected animals. Furthermore, if left untreated, *Salmonella* disseminated throughout the body of TH mice by day 5 post infection (Figure S3A); however, butyrate treatment effectively limited pathogen spread and significantly reduced the overall bioluminescence intensity (Figure S3B). Importantly, butyrate supplementation significantly reduced the bacterial burden in the intestines (cecum and colon) as well as in mesenteric lymph node (MLN), spleen, and liver (Figure 5C). Finally, tributyrin treatment also resulted in a slight reduction of STM-induced inflammation in the cecum compared to the untreated STM-infected animals (Figure S5B).

**Figure 5.**
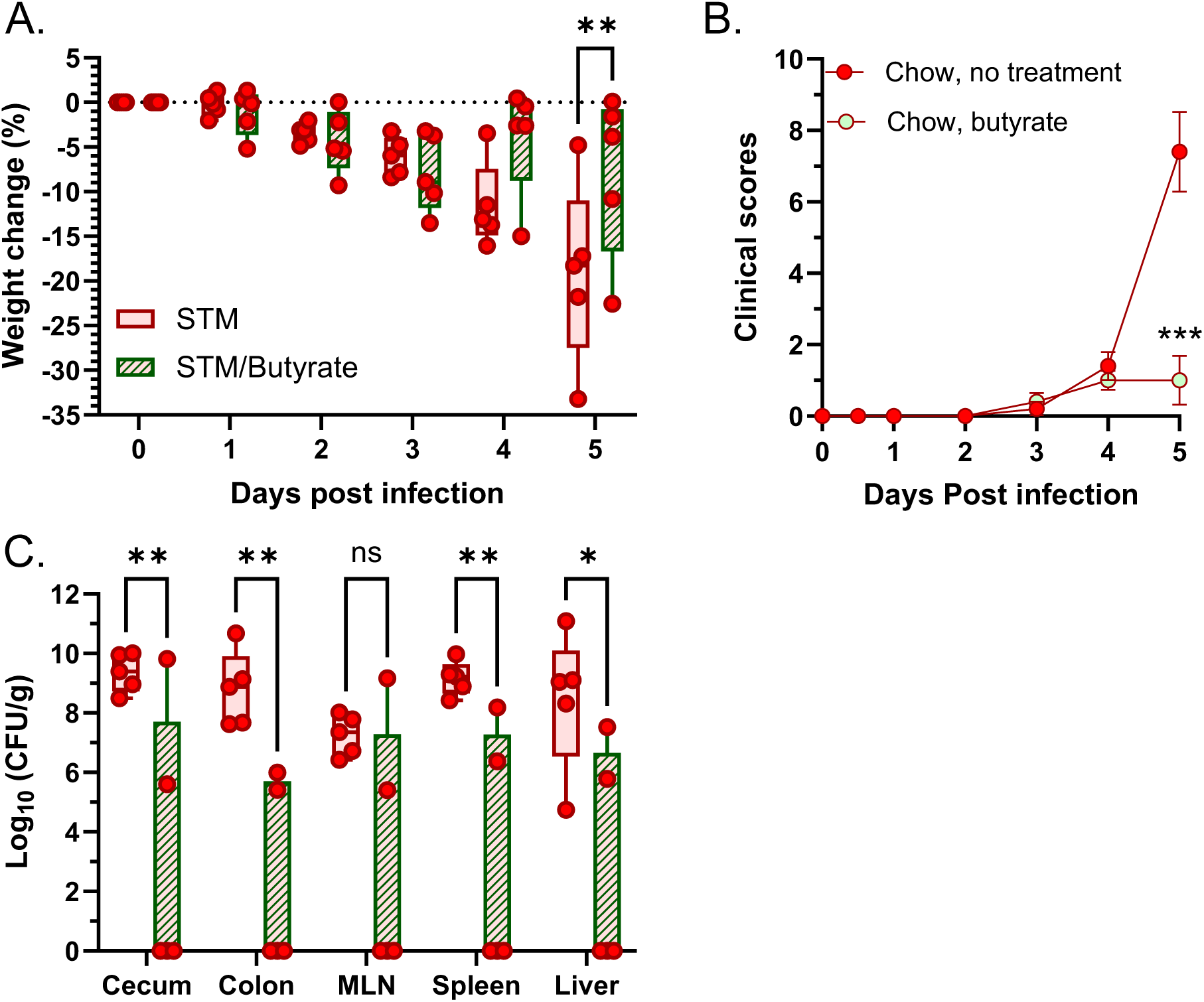
Butyrate supplementation limits *Salmonella* colonization in normoglycemic mice on chow diet. **A.** Body weight changes in response to STM infection followed by tributyrin treatment compared to untreated STM-infected mice. **B.** Clinical scores post infection. **C.** Butyrate supplementation reduces STM burden in the gut (cecum, colon), mesenteric lymph node (MLN), spleen and liver at day 5 post infection compared to the untreated STM-infected animals. Asterisk indicates significance in two-way ANOVA with Šìdák’s multiple comparison’s test, *p*<0.05.

Next, we orally infected isogenic hyperglycemic HFD mice with the same number of bacteria as was used for infection of normoglycemic animals followed by a daily tributyrin treatment. Unexpectedly, STM-infected mice treated with tributyrin experienced an accelerated weight loss 24 hours post infection (Figure 6A). Also, butyrate supplementation did not improve the overall clinical scores compared to the untreated infected mice (Figure 6B) and all infected animals were euthanized on day 4 post infection due to severity of the disease. Similarly, tributyrin treatment did not change bioluminescence intensity (Figure S4), indicating the inability of butyrate supplementation to limit *Salmonella*’s spread in a hyperglycemic host. In accordance with previous observations, the levels of bacterial colonization in the intestines and in the extraintestinal tissue (MLN, spleen and liver) were not significantly different between samples collected from the infected animals with or without tributyrin treatment (Figure 6C).

**Figure 6.**
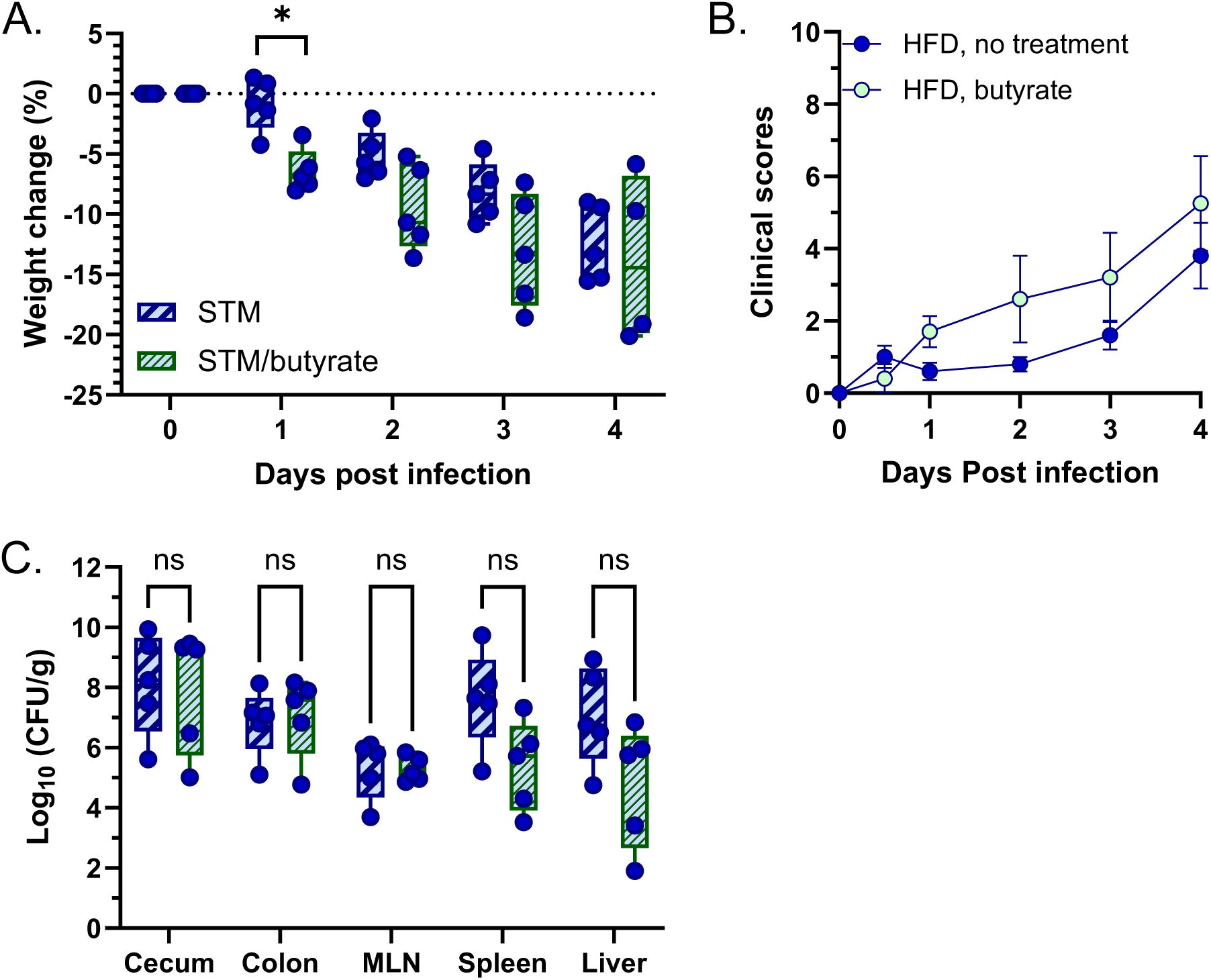
Butyrate supplementation fails to limit *Salmonella* colonization in hyperglycemic HFD mice. **A.** Body weight changes in response to STM infection followed by tributyrin treatment compared to untreated STM-infected mice. **B.** Clinical scores post infection. **C.** Butyrate supplementation does not affect STM burden in the gut (cecum, colon), mesenteric lymph node (MLN), spleen and liver at day 4 post infection compared to the untreated STM-infected animals. Asterisk indicates significance in two-way ANOVA with Šìdák’s multiple comparison’s test, *p*<0.05.

This surprising failure of the butyrate supplementation in limiting *Salmonella* colonization in hyperglycemic animals may be related to preexisting mild inflammation caused by HFD. Butyrate can be damaging to epithelial cells in the pro-inflammatory environment (63). Quantification of luminal TNFα levels in the cecum of uninfected mice confirmed that the animals in HFD had increased amounts of this pro-inflammatory cytokine compared to mice in chow-fed group regardless of glucose levels in their blood (Figure S5A). Because *Salmonella* itself also triggers production of TNF-α by the host (64), the combination of tributyrin intervention with infection resulted in detectable epithelial damage in the cecum of STM-infected/butyrate treated mice compared to the untreated infected animals (Figure S5C).

## Discussion

The current understanding of *Salmonella enterica* serovar Typhimurium pathogenesis is based on the use of animal models developed to provide critical insights into different aspects of disease caused by this prevalent pathogen. For instance, the use of calves has allowed researchers to study signs of disease that are parallel to the symptoms observed in humans (65) while pre-treatment of mice with streptomycin has supported studies related to *Salmonella*-induced inflammation (66). The use of mice with intact microbiota has advanced studies focused on the mechanisms of transmission and virulence factors needed for pathogen’s colonization (67). However, the *Salmonella* research up to date has been largely centered on healthy hosts, although a substantial proportion of the human population is affected by obesity, type 2 diabetes, and other metabolic diseases. The studies of *Salmonella* pathogenesis in the context of metabolic disease have been delayed due to the lack of an animal model that can accurately replicate these chronic conditions.

TALLYHO mice, an established inbred polygenic model of type 2 diabetes with moderate obesity (68), provide the opportunity to interrogate previously neglected aspects of *Salmonella* disease in the context of diabesity. Although polygenic models offer a more accurate model of human chronic condition compared to monogenic models, there is no proper wild type control available. For instance, SWR/J is a recommended control strain for TALLYHO mice by the Jackson Laboratory; however, these strains share only 86.8% of their genotype and 67.1% of their haplotype. As a result, many aspects of metabolic and skeletal characteristics are different between these mouse strains regardless of blood glucose status (69). Therefore, normoglycemic TH males are now considered to be a more appropriate strain-matched non-diabetic control (34, 69).

While both males and females develop obesity with age, hyperglycemia is limited to TH males only (68). This closely resembles the situation with male predominance of type 2 diabetes in human population at younger age (70, 71). Furthermore, it was shown that ovariectomy in TH female mice promotes chronic hyperglycemia, which supports a protective role of estrogen in diabetes development (72). Importantly, in TH mice, chronic hyperglycemia gradually develops by 16 weeks of age as opposed to up to six months of HFD exposure in C57BL/6J mice required for hyperglycemia development in this alternative model of diabesity (31).

Hyperglycemia development in male TH mice can be promoted by diet with high sucrose or high fat content (35, 73). In agreement with earlier publications, feeding TH mice with HFD (45% fat) for 8 weeks resulted in development of chronic hyperglycemia in the majority of animals (Figure 1B). Nevertheless, a small fraction of animals in this group remained normoglycemic and served as an additional control for diet in the infection studies.

In accordance with previous reports (74, 75), consumption of HFD did not influence spleen mass in normoglycemic mice compared to the control chow-fed animals. Nevertheless, persistent hyperglycemia resulted in significant splenomegaly in chow-fed TH mice (Figure S1). The significant enlargement of liver and kidney in hyperglycemic animals is consistent with previously noticed association of diabesity with low-grade inflammation known as metaflammation (76).

To date, the use of TALLYHO mice in host-pathogen interaction research has been limited to infection studies focused on wound and pressure ulcer healing, as well as periodontitis, conditions that are common in diabetic patients (77–80). However, because TALLYHO/JngJ, BALB/c and C57BL mouse strains share the same *Slc11a1* haplotype, one can predict that TH mice would be susceptible to *Salmonella* infection. Indeed, oral infection of chow-fed or HFD normoglycemic male TH mice with 10^6^ CFU of *Salmonella* Typhimurium strain resulted in robust colonization of the gut followed by systemic spread later in infection (Figures 2, 5 and S3). Unexpectedly, the colonization pattern was drastically different in HFD animals with hyperglycemia and led to an accelerated escape from the gut and overall increase in clinical scores (Figure 2). These findings are congruent with reports of salmonellosis in diabetic patients, which is associated with developing of focal metastatic infections (11–19, 21, 22).

The increased susceptibility of patients with type 2 diabetes to salmonellosis was originally linked to reduced acidity in the stomach and impaired intestinal motility (23). However, persistent hyperglycemia can also lead to increased translocations of bacterial LPS from the intestine to the bloodstream in diabetic patients (53, 81–84). Accordingly, we detected higher levels of LPS in the serum of HFD mice with hyperglycemia compared to normoglycemic animals on the same diet (Figure 3A).

Type 2 diabetes results in a reduction of butyrate-producing bacteria in the gut. This phenomenon was first noticed in diabetic patients (36–40), and was later confirmed in rodents (85). Intestinal short chain fatty acids (SCFA) are among the major products of fiber fermentation in the gut. A typical chow diet is rich in insoluble and soluble fibers which can be metabolized by microbiota. On the contrary, refined diets, including diets with high fat content, are low in fiber and contain only cellulose powder which is insoluble and non-fermentable (86). As a result, production of SCFA is significantly reduced in the intestines of mice fed refined diets regardless of the dietary fat content compared to the chow-fed animals (87). Similarly, SCFA concentrations were lower in the cecal samples collected from TH mice in HFD groups regardless of their blood glucose status compared to the animals maintained on the chow diet. However, the amount of cecal SCFA, including butyrate, was substantially reduced in chow TH mice with persistent hyperglycemia compared to the normoglycemic animals despite consumption of food rich in fiber (Figures 3 and S2).

Many studies have demonstrated that butyrate can act as an anti-inflammatory agent. For instance, butyrate reduces gut mucosal inflammation through inhibition of NF-κB transcription factor (88) that regulates expression of proinflammatory cytokines (42). Similarly, butyrate lowers the level of proinflammatory cytokines IL-1β and TNF-α, which are secreted by peripheral blood mononuclear cells (PBMC) in response to *Mycobacterium tuberculosis* (Mtb) cell lysate (89). These cytokines are known to be elevated in peripheral blood of type 2 diabetes patients (90). Butyrate also activates Peroxisome-Proliferator-Activated-receptor γ (PPARγ) (91, 92) and inhibits INF-γ signaling (42).

Butyrate is also known to repair and enforce the barrier function of intestinal epithelial cells (42). However, the effect of butyrate on the gut barrier function appears to be concentration dependent. While butyrate promotes intestinal integrity at low concentrations, it can damage barrier function by inducing apoptosis at higher concentrations (42). The amount of tributyrin used in our experiments is comparable to previous studies (93, 94) and, as expected, butyrate supplementation reduced LPS leakage from the intestines in both normoglycemic TH mice on chow diet and hyperglycemic HFD animals (Figure 4). Furthermore, butyrate effectively limited *Salmonella* escape from the gut in normoglycemic animals (Figure 5 and S3) but paradoxically failed to limit pathogen spread in hyperglycemic HFD mice (Figure 6 and S4).

Notably, butyrate can be detrimental for intestinal barrier integrity in the presence of inflammatory mediators (TNF-α and INF-γ) (63). Interestingly, our data show that TNF-α levels are elevated in the cecal lumen of HFD mice prior to infection (Figure S5A). It is possible that because infections with *Salmonella* also lead to increased production of TNF-α (64), butyrate treatment promoted epithelial damage in STM-infected hyperglycemic mice (Figure S5) through activation of apoptosis-related proteins (63). It was also proposed that SCFA, including butyrate, can be differentially handled in obese insulin-resistant state as opposed to the healthy host and that may contribute to the unexpected lack of butyrate therapy benefits in patients with metabolic syndrome (95). While this phenomenon needs to be investigated further, since several clinical trials with butyrate to treat chronic conditions are underway, it is important to make clinicians aware of potential risks and benefits associated with this metabolite.

In conclusion, our study established a model to study bacterial pathogenesis in the context of diabesity; we demonstrated that the increased susceptibility of hyperglycemic mice to *Salmonella* infection is at least in part due to increased intestinal permeability and illustrated that the strategies appropriate for control of infection in the healthy host might not be suitable for pathogen containment in the patients with chronic conditions. Further elucidation of the underlying reasons for butyrate failure in limiting *Salmonella* colonization and extraintestinal spread in the diabetic host will significantly expand the current understanding of key metabolic alterations and physiological changes in the gastrointestinal tract of patients with type 2 diabetes and will provide clues to therapeutically counterbalancing these effects.

## Materials and Methods

### Bacterial strains, media, and growth

A fully virulent nalidixic acid resistant derivative of bioluminescent *Salmonella enterica* ser. Typhimurium strain was created by moving *hisG*::*rpsM-luxCDABE* (96) into a spontaneously nalidixic acid-resistant derivative of ATCC 14028 (97) using P22 transduction (98). The resulting strain, LB62, was tested for light production using IVIS Lumina XRMS system (Perkin Elmer). Bacterial cultures were grown in Luria-Bertani (LB) broth (Difco) supplemented with 50 mg/liter nalidixic acid (Chem-Impex) at 37°C with agitation (200 rpm).

### Animals

All procedures described in this study were approved by the Marshall University Animal Care and Use Committee. Seven to eight weeks old male TALLYHO/JngJ mice (Jackson Laboratories, Bar Harbor, Maine) were used throughout the study. TALLYHO/JngJ (TH) inbred mice are a polygenetic model for non-insulin dependent type 2 diabetes mellitus (T2D) where the development of hyperglycemia is limited to male mice (68). After one-week acclimation to Marshall Animal Research Facility, animals were randomly assigned to one of the two diets for eight weeks: a standard chow (5001 Rodent diet, LabDiet) or a gamma-irradiated 45% high fat diet (D12451i, Research Diets) and were monitored for hyperglycemia development. Mice were housed in groups of three with *ad libitum* access to sterile food and water in a temperature and humidity-controlled room with a 12-hour light/dark cycle. The body weights were taken weekly and used to track weight gain. The non-fasting blood glucose levels were measured in all TH mice at 10-, 12-, 14-, and 16-weeks of age using a OneTouch Ultra glucometer (99). The maximal level of blood glucose that can be determined with this glucometer is 600 mg/dl. Mice with average glucose levels >300 mg/dl were considered hyperglycemic (28).

### Oral infection with bioluminescent *Salmonella*

Five 16-week-old TH mice with persistent hyperglycemia from HFD group and five age matched TH mice with normal blood glucose levels from chow group were infected by gavage with 1x10^6^ bioluminescent bacteria in 100 µl of sterile phosphate-buffered saline (PBS). The inoculum was grown at 37°C with aeration, serially diluted and plated for CFU (colony forming units) enumeration to determine the exact titer. Additional normoglycemic HFD animals were infected in a similar fashion and served as additional control for the diet contribution to the infection outcome. Post-infection, animals were imaged daily using IVIS Lumina XRMS system (Perkin Elmer) to follow the pathogen spread. Uninfected animals with or without hyperglycemia from HFD or chow groups were imaged in parallel to estimate the background signal. For imaging, mice were anesthetized in an oxygen-rich induction chamber with 3% isoflurane. Animals were imaged in the ventral position; anesthesia was maintained during the entire imaging session by using a nose cone 1.5% isoflurane-oxygen delivery device in the imaging chamber. Data acquisition and image analysis was performed using Living Image 4.7.3 software (Perkin Elmer). Bioluminescence was measured as radiance (photons/s/cm^2^/sr). To allow comparison between images from different days, a background cutoff equal to 10^6^ photons/s/cm^2^/sr was applied to all images. For quantification of the detected bioluminescence over the course of infection, the body of the mouse was selected using ROI (region of interest) tool and the total radiance was recorded.

Animals were monitored for weight loss and signs of distress and were humanely euthanized on day 5 post infection or when the cumulative clinical score is equal to 5 or higher. Clinical signs were scored based on the following criteria (on a scale from 0 to 3 each): body conditioning (0 - well-conditioned; 1 - animal lost between 10-15% body weight; 2 – animal lost between 15-20% body weight; and 3 - animal lost >20% body weight), physical appearance (0 – normal; 1 – lack of grooming; 2 – squinting and hunching, 3 – excessive hunching and squinting), unprovoked and provoked behavior (0 – normal; 1 - minor changes; 2 – moderate changes; 3 – severe changes). The spleens, livers, kidneys, mesenteric lymph nodes, ceca and colons were collected in individual tubes containing sterile PBS, weighed, homogenized, serially diluted, and plated on nalidixic acid containing LB plates for CFU enumeration.

### Analysis of intestinal permeability

To measure the potential impact of persistent hyperglycemia on gut barrier function, serum was collected from 16-week-old TH mice with and without hyperglycemia after 8 weeks of chow or HFD exposure. Circulating LPS levels were determined using the PyroGene Recombinant Factor C Endpoint Endotoxin Fluorescent assay (Lonza) following manufacturer’s recommendations. Serum samples from uninfected mice were diluted in LAL Reagent water (1:1,000 – 1:10,000) and plated in triplicate in a sterile 96 well plate. The plate was read at 0 and 60 minutes with results corrected to 0 minutes (excitation at 360nm and emission at 460nm; BioTek Synergy HTX). A standard curve was calculated from the supplied standards with a sensitivity of 0.0005 EU/mL.

### Gas Chromatography-Mass Spectrometry analysis

For short chain fatty acid profile analysis, cecal contents from uninfected animals from HFD and chow groups were collected, resuspended in sterile PBS, weighed, and placed on ice. Samples were processed using an established protocol (93, 100, 101). Briefly, samples were vortexed for 2 min and separated by centrifugation at 13,000 rpm for 15 min at 4°C. Supernatants were transferred to a new tube, mixed with deuterated acetate, propionate, and butyrate (CDN Isotopes) as the internal standard (5 µM each), and dried using a SpeedVac concentrator. Samples were dissolved in pyridine in 1:1 ratio, sonicated for 1 minute, and incubated at 80°C for 20 min. For derivatization, an equal amount of *N-tert*-butyldimethylsilyl-*N*-methyltrifluoroacetamide with 1% *t*-BDMCS (*tert*-butyldimethylchlorosilane, Cerilliant) was added, and the samples were incubated for 1 h at 80°C. After centrifugation at 14,000 rpm for 1 min, samples were transferred to autosampler vials for GS-MS analysis (Shimadzu, TQ8040) using a Rtx-5Sil MS column (30 m × 0.25 mm × 0.25 μm; Shimadzu). The injection temperature was 250°C, and the injection split ratio was set to 1:100 with an injection volume of 1 μL. The oven temperature started at 50°C for 2 min, increasing to 100°C at 20°C per minute and to 330°C at 40°C per min, with a final hold at this temperature for 3 min. The flow rate of the helium carrier gas was kept constant at a linear velocity of 50 cm/s. The interface temperature was 300°C. The electron impact ion source temperature was 200°C, with a 70-V ionization voltage and a 150-μA current. To measure acetate, propionate and butyrate, single ion monitoring was used with the following target and reference (italicized) ions: acetate – m/z 117, *75, 159*; propionate - m/z 131, *75, 132*; butyrate - m/z 145, *146, 75*; butyrate-d_7_ - m/z 153, *76, 152*, respectively.

### Tributyrin treatment

Five 16-week-old TH mice with persistent hyperglycemia from HFD group and five age matched TH mice with normal blood glucose levels from chow group were infected by gavage with 1x10^6^ bioluminescent bacteria in 100 µl of sterile phosphate-buffered saline (PBS) as described above. Mice were treated daily with 100 µl of undiluted tributyrin (TCI) (93) delivered by gavage starting on the day of infection. Uninfected HFD animals with hyperglycemia and normoglycemic mice from the chow group were treated with tributyrin in parallel. Animals were imaged daily using IVIS Lumina XRMS system (Perkin Elmer) to follow the pathogen spread. Following euthanasia, the spleens, livers, mesenteric lymph nodes, ceca and colons were collected in individual tubes containing sterile PBS, weighed, homogenized, serially diluted, and plated on nalidixic acid containing LB plates for CFU enumeration.

### ELISA for TNF-**α** measurement

To measure TNF-α levels, cecal contents from uninfected normo- and hyperglycemic mice from chow and HFD groups were diluted to 100 mg/ml in sterile PBS. Total protein was measured for each sample using a BCA assay (Millipore). TNF-α was measured using mouse TNF-α DuoSet ELISA kit (R&D Systems) per manufacturer’s instructions, reading at 450nm with correction from 570nm and pathlength. Three biological replicates per condition with three technical replicates per mouse were used. Results were normalized to total protein per sample and expressed as pg TNF-α / mg total protein.

### Histopathology analysis

The tissue was fixed in 10% phosphate buffered formalin. Samples were paraffin embedded, cut, and stained with hematoxylin and eosin at the Cabell Huntington Hospital Histology lab. Blinded slides were scored by Board-certified pathologist for signs of inflammation using criteria described in (102). The combined score between 0- 5 corresponded to normal to mild inflammation; the score greater than 5 indicated moderate inflammation.

## Data analysis

Statistical analyses were performed using GraphPad Prism v.10.6.1.

## Acknowledgments

This work was supported by the WV-INBRE Chronic Disease Research Program (NIH award P20GM103434 to the West Virginia IDeA Network for Biomedical Research Excellence) and by a grant from the West Virginia Clinical and Translational Science Institute (NIH U54GM104942). We also sincerely thank Julia Cardot for the valuable discussions that contributed to this research.

## Supplementary Figure legends

**Figure S1. Persistent hyperglycemia leads to enlargement of spleen, liver and kidney.** Asterisk indicates significance in multiple Mann-Whitney U test with Holm- Šìdák method, *p*<0.05

**Figure S2. Persistent hyperglycemia and HFD lead to reduction in the total cecal SCFA.** Asterisk indicates significance in multiple unpaired t test with Welch correction test using Holm-Šìdák method, *p*<0.05.

**Figure S3. Butyrate supplementation limits *Salmonella* spread in normoglycemic mice on chow diet. A.** 16-weeks-old male TH mice maintained on chow diet were orally infected with 1x10^6^ CFU of *Salmonella* strain expressing chromosomally encoded *hisG::rpsM-luxCDABE,* treated with daily oral dose of tributyrin (as in Figure 4), and imaged using IVIS Lumina XRMS III system at day 5 post infection; **B.** Bioluminescence quantification was done using Living Image software. Asterisk indicates significance in 2-way ANOVA with Šìdák’s multiple comparisons test, *p*<0.05

**Figure S4. Butyrate supplementation fails to limit *Salmonella* spread in hyperglycemic HFD mice. A.** 16-weeks-old male TH mice maintained on HFD were orally infected with 10^6^ CFU of *Salmonella* strain expressing chromosomally encoded *hisG::rpsM-luxCDABE,* treated with daily oral dose of tributyrin (as in Figure 4), and imaged using IVIS Lumina XRMS III system; **B.** Bioluminescence quantification was done using Living Image software. Data were analyzed using 2-way ANOVA with Šìdák’s multiple comparisons test.

**Figure S5. Butyrate is detrimental for intestinal barrier integrity of STM-infected animals in the presence of inflammatory mediator TNF-**α**. A.** Cecal TNF-α levels were measured using ELISA. Asterisk indicates significance in multiple unpaired t test with Welch correction Holm-Šìdák method, *p*<0.05. **B, C.** Sections of cecum were collected from normoglycemic TH mice at day 5 post infection (B), or from TH mice at day 4 post infection (C); fixed in formalin, paraffin embedded, cut, and stained with hematoxylin and eosin. Stained sections were scored for the signs of inflammation. A combined score of <5 corresponds to normal or mild inflammation; a combined score of >5 indicates moderate to severe inflammation.

