## Supplementary figures and images for "Type 2 diabetes increases susceptibility to invasive *Salmonella* Typhimurium despite butyrate supplementation"

### Figure S1

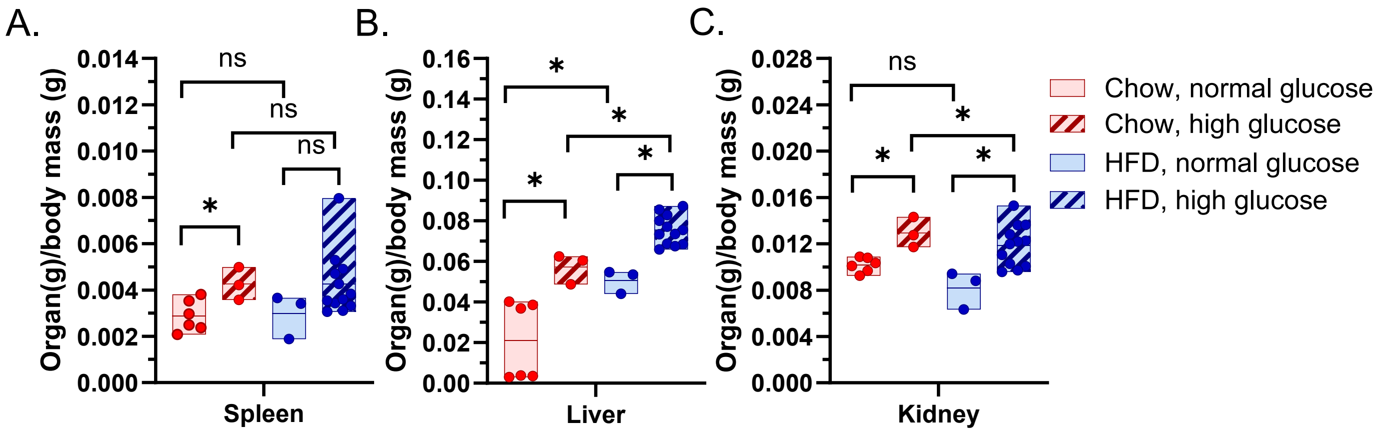

### Figure S2

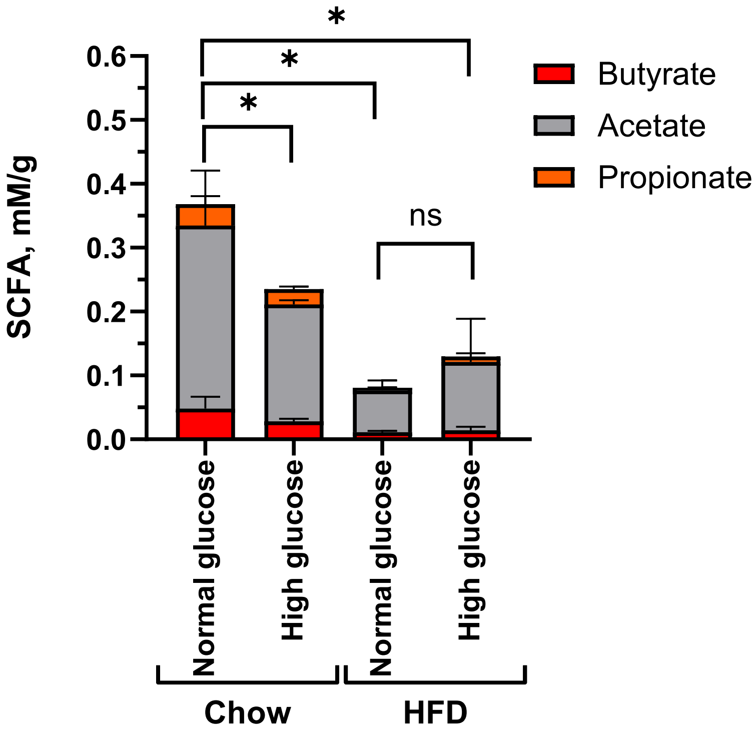

### Figure S3

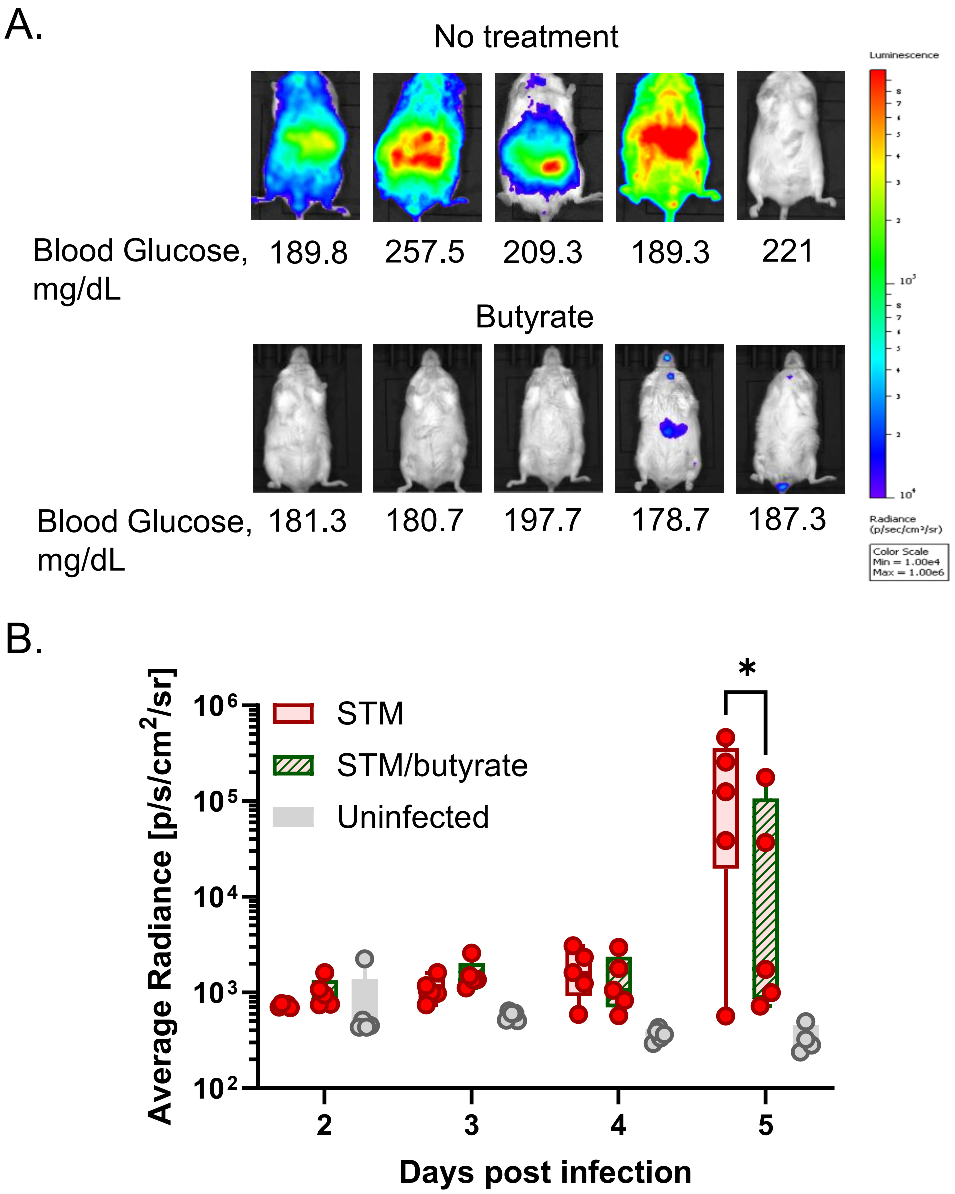

### Figure S4

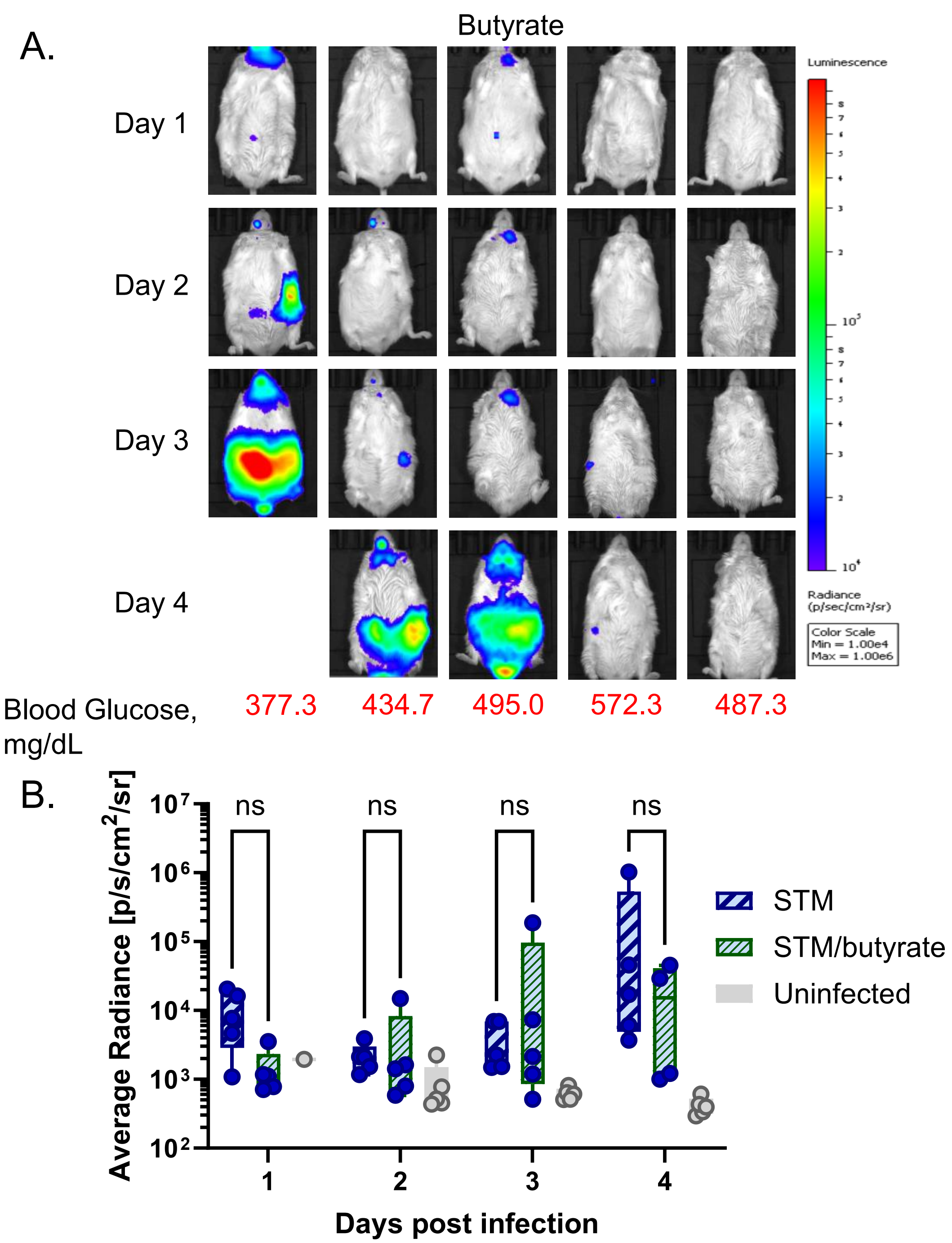

### Figure S5

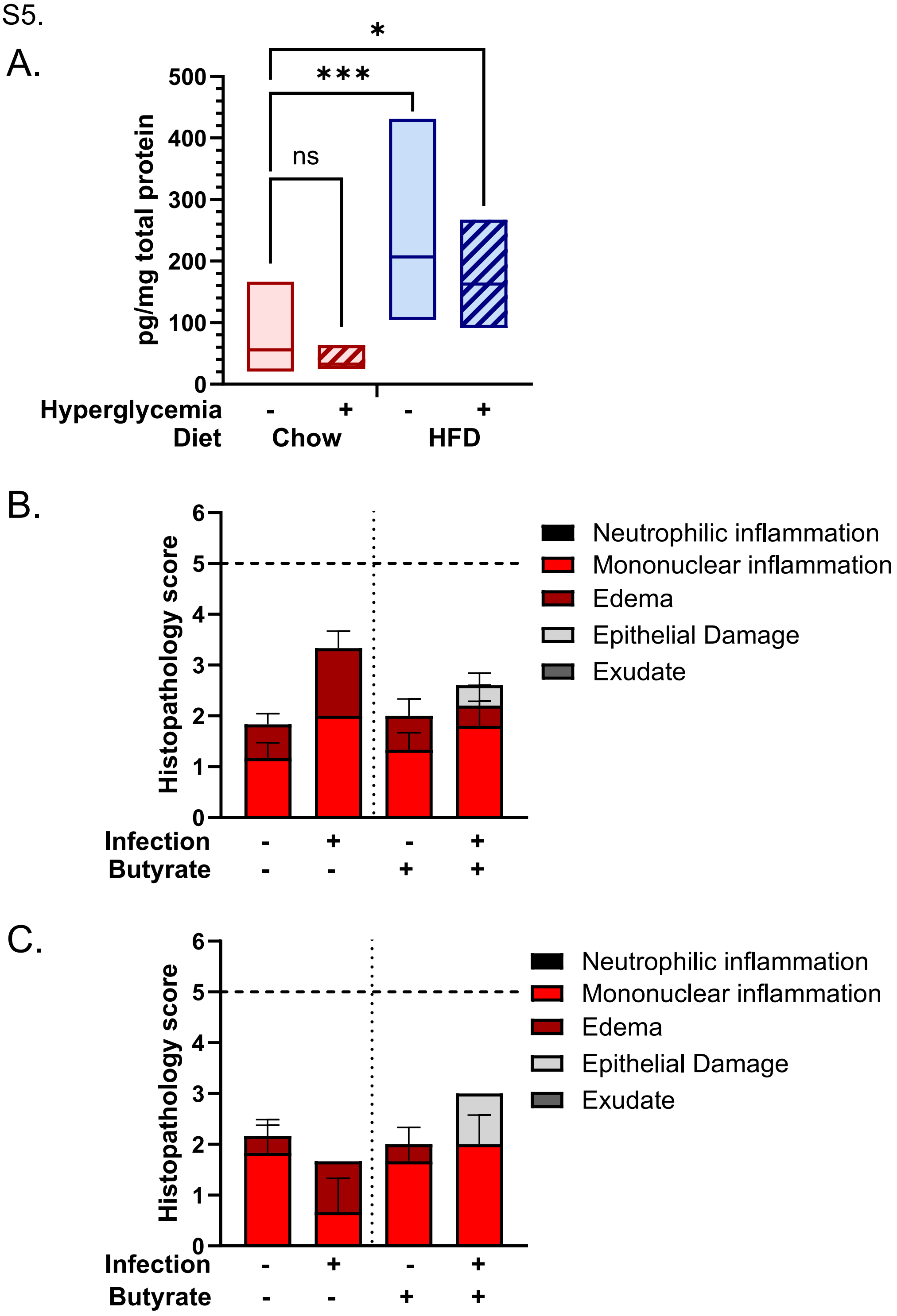
